# Modeling noise mechanisms in neuronal synaptic transmission

**DOI:** 10.1101/119537

**Authors:** Abhyudai Singh

**Affiliations:** Department of Electrical and Computer Engineering Department of Biomedical Engineering and Department of Mathematical Sciences University of Delaware, Newark, DE, USA

## Abstract

In the nervous system, communication occurs via synaptic transmission where signaling molecules (neurotransmitters) are released by the presynaptic neuron, and they influence electrical activity of another neuron (postsynaptic neuron). The inherent probabilistic release of neurotransmitters is a significant source of noise that critically impacts the timing of spikes (action potential) in the postsynaptic neuron. We develop a stochastic model that incorporates noise mechanisms in synaptic transmission, such as, random docking of neurotransmitter-filled vesicle to a finite number of docking sites, with each site having a probability of vesicle release upon arrival of an action potential. This random, burst-like release of neurotransmitters serves as an input to an integrate-and-fire model, where spikes in the postsynaptic neuron are triggered when its membrane potential reaches a critical threshold for the first time. We derive novel analytical results for the probability distribution function of spike timing, and systematically investigate how underlying model parameters and noise processes regulate variability in the inter-spike times. Interestingly, in some parameter regimes, independent arrivals of action potentials in the presynaptic neuron generate strong dependencies in the spike timing of the postsynaptic neuron. Finally, we argue that probabilistic release of neurotransmitters is not only a source of disturbance, but plays a beneficial role in synaptic information processing.

## I. INTRODUCTION

Communication between neurons occurs through chemical synapses, where signals in the form of neurotransmitters are passed from a presynaptic neuron to a postsynaptic neuron. Synaptic structures are generally present at the axon terminals of the presynaptic neuron, and consist of neurotransmitter-filled vesicles that are loaded on to a finite number of docking sites. An action potential reaching the axon terminal triggers opening of voltage-gated calcium channels. Calcium influx into the axon terminal causes fusion of docked synaptic vesicles to the cell membrane, and release of neurotransmitters. The depletion of docked synaptic vesicles leads to the system becoming less responsive to the next action potential, creating some sort of memory that is often referred to as synaptic depression [1]. Released neurotransmitters bind to receptors on the postsynaptic neuron, and open ion channels that drive short-term changes in the membrane potential of the postsynaptic neuron. While individual neurons are known to receive synaptic contacts (both inhibitory and excitatory) from thousand of other neurons, we here consider the simplest case of a single presynaptic neuron forming an excitatory synapse with a single postsynaptic neuron

Much prior work treats neurotransmitter release as a deterministic processes [2–6], but these models fail to capture the variability introduced at each trial by the inherent probabilistic nature of vesicle release and other noise mechanisms [7–10]. Moreover, several experimental and computational works have argued that these stochastic effects are functionally important for understanding information flow across a synapse [11–16]. As part of this contribution we consider a stochastic model of synaptic transmission that incorporates three different noise mechanisms:

1. Random arrival of action potentials at the axon terminal, with inter-arrival times drawn from an arbitrary probability distribution.
2. When the action potential arrives, each docking site loaded with a vesicle has a probability of releasing the vesicle.
3. Empty sites recover probabilistically, and over time they become occupied by synaptic vesicles.

Before formally introducing the stochastic model and our main analysis, we discuss some mathematical preliminaries.

## II. MATHEMATICAL PRELIMINARIES

We start by recalling some basic probability concepts related to the binomial distribution, as this distribution arises ubiquitously in the modeling of noisy synaptic transmission. If the random variable *X* follows the binomial distribution with parameters *M* ∈ {1, 2,…} (the number of trials) and *p* ∈ [0-1] (success probability in each trial), then we write

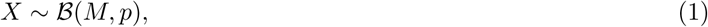
and its probability mass function is given by

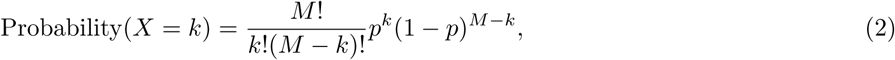
*k* ∈ {0,1, 2,…}. The mean and variance of *X* is 
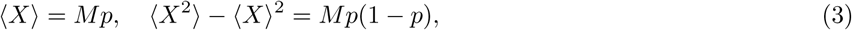
respectively, where 〈 〉 denotes the expected value operator. Moreover, 
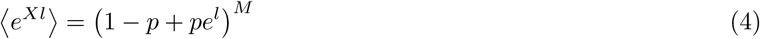
denotes the moment generating function (MGF) of *X*. A key result that will be used later relates to conditional binomials. If 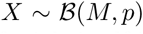, and we define a new random variable *Y* such that conditioned on 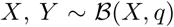 (i.e., the number of trials itself is binomially distributed) then 
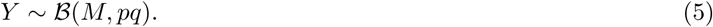

This result can be can be straightforwardly proven using the MGF in (4) and we omit the proof due to space constraints. Next, we describe a stochastic model for neurotransmitter release from the presynaptic neuron, and show how binomial concepts introduced here aid the model analysis.

## III. STOCHASTIC MODELING OF NEUROTRANSMITTER RELEASE

Let action potentials arrive at the presynaptic neuron at discrete times *t_i_, i* ∈ {1, 2,…}, and the intervals

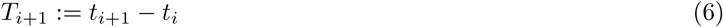
be independent and identically distributed (i.i.d.) random variables drawn from a given probability density function. We denote by 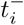 and 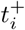, the time just before and after the *i^th^* action potential, respectively. Upon arrival of an action potential, neurotransmitter molecules are released based on a stochastic model with the following three ingredients:

1. There are *M* docking sites on the presynaptic neuron, and each site can either be empty, or occupied by a synaptic vesicle that is filled with neurotransmitters.
2. When the action potential arrives, each occupied site has a probability *p_r_* ∈ [0-1] of releasing the vesicle and becoming empty. The release process is assumed to occur instantaneously in time.
3. In the time interval between two successive action potentials, an occupied sites remains occupied, and an empty sites become occupied with a rate *k*.

As a consequence of the last point, if a site is empty after the *i^th^* action potential, then the probability *p_i_* ∈ [0-1] of a vesicle docking there before the arrival of the *i* + 1*^th^* action potential is 
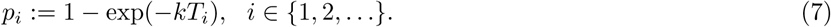

Intuitively, if the next action potential takes much longer to arrive by random chance, then there is a higher probability of an empty site getting occupied by a synaptic vesicle. Note that the probabilities *p_i_* are themselves i.i.d random variables since *T_i_* are i.i.d. As an example, if the action potential arrives based on a Poisson process and *T_i_* is exponentially distributed with mean 〈*T_i_*〉, then as per (7) the probability density function of *p_i_* is given by 
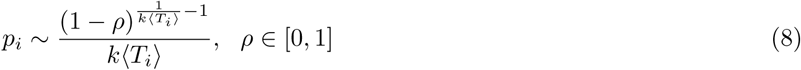
with the following mean and variance

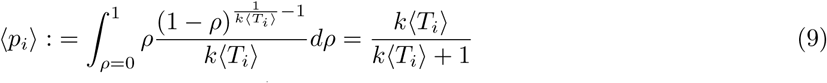

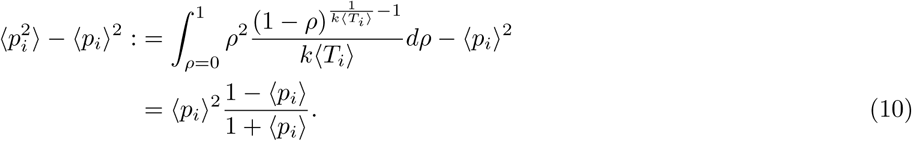

As expected, 〈*p_i_*〉 increases with increasing gap between successive action potentials, and in the limit 〈*T_i_*〉 →∞, *p_i_* = 1 with probability one. Another scenario that we will discuss later on is deterministic arrival of action potentials, for which *T_i_* = 〈*T_i_*〉 with probability one, and 
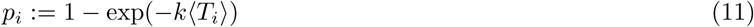
with probability one. A sample path of the number of synaptic vesicles released over time is shown in Fig. 1. For convenience, notations used in the model and subsequent analysis are summarized in Table I. The following Theorem quantifies the amount of neurotransmitter released.

**FIG. 1.**
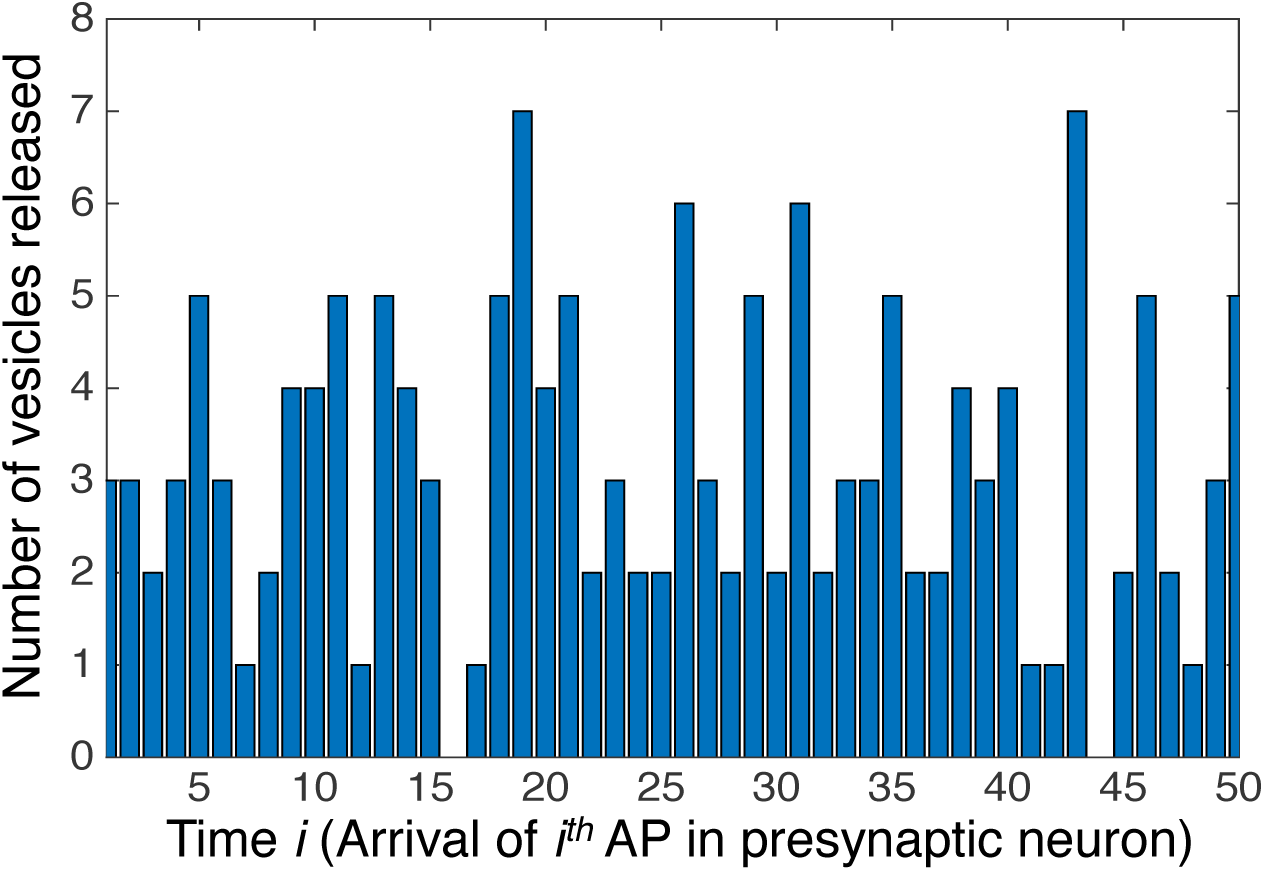
A sample realization of the number of synaptic vesicles released each time an action potential (AP) arrives at the presynaptic neuron. For this plot, the number of docking site is taken as *M* = 9, with each occupied site having a probability of release *p_r_* = 0.5. Action potentials are assumed to arrive at deterministic time intervals, and an empty site has a probability *p_i_* = 0.5 of getting occupied before the arrival of the next action potential.

**TABLE I.** Summary of Notations

|  |  |
| --- | --- |
| AP | Action potential |
| $t_i$ | Time of arrival of the $i^{th}$ AP in presynaptic neuron |
| $t_i^-$ | Time just before $i^{th}$ AP in presynaptic neuron |
| $t_i^+$ | Time just after $i^{th}$ AP in presynaptic neuron |
| $T_i$ | Random variable denoting inter-arrival times $t_{i+1} - t_i$ |
| $\langle T_i \rangle$ | Mean inter-arrival time of AP |
| Input frequency | $1/\langle T_i \rangle$ |
| $M$ | Number of docking sites for synaptic vesicles |
| $p_r$ | Probability of vesicle release per occupied site |
| $k$ | Rate at which an empty site becomes occupied |
| $p_i$ | Probability of an empty site becoming occupied in the time interval $T_i$ |
| $N_i^-$ | Number of occupied sites just before $i^{th}$ AP |
| $N_i^+$ | Number of occupied sites just after $i^{th}$ AP |
| $Z_i$ | Number of vesicles released when $i^{th}$ AP arrives |
| $CV_Z$ | Coefficient of variation (stdev/mean) of $Z_i$ |
| $\tilde{p}_i$ | Probability of a site being occupied at $t_i^-$ |
| $c$ | Number of neurotransmitters per vesicle |
| $\gamma_z$ | Decay rate of neurotransmitter molecules |
| $z(t)$ | Neurotransmitter count in the synaptic gap |
| $\langle z \rangle$ | Mean neurotransmitter count at steady-state |
| $CV_z$ | Coefficient of variation (stdev/mean) of $z(t)$ |
| $v(t)$ | Membrane potential of the postsynaptic neuron |
| $CV_v$ | Coefficient of variation (stdev/mean) of $v(t)$ |
| $\tau_v$ | Membrane time constant |
| $v_{th}$ | Threshold for AP firing in the postsynaptic neuron |
| $T^*$ | Time to next AP in the postsynaptic neuron |
| Output frequency | $1/\langle T^* \rangle$ |
| $CV_{T^*}$ | The coefficient of variation (stdev/mean) of $T^*$ |

### Theorem 1

Let *c* denote the number of neurotransmitters in each synaptic vesicle. Then, upon arrival of the i^th^ action potential, the number of neurotransmitter molecules released is *cZ_i_*, where

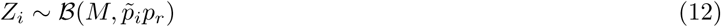

is the number of synaptic vesicles released. The probability 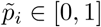 in (12) follows the random discrete-time system

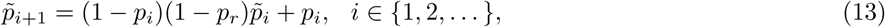

where *p_i_* are i.i.d random variables defined in (7).

**Proof** Let 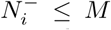 denote the number of occupied sites at time 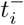 (just before the arrival of the *i^th^* action potential). Given the probability of vesicle release, *p_r_*, the number of occupied sites releasing vesicles is 
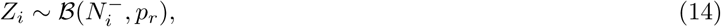
and the number of occupied sites at time 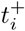 is given by 
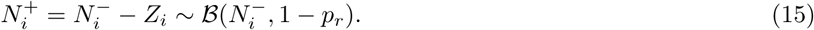

Now the number of empty sites at 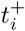 is 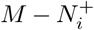, and each empty site has a probability *p_i_* of getting occupied before the arrival of *i* + 1*^th^* action potential. Then, the number of occupied sites just before the arrival of the *i* + 1*^th^* action potential is 
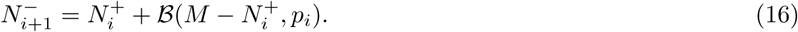

To solve this stochastic map we assume that 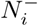 follows a binomial distribution 
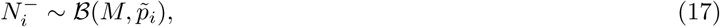
and apply the concept of conditional binomials from (5) to (15) and (17) 
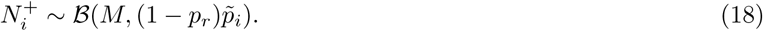

In the Appendix A, we show using (18) that the right-hand-side of (16) follows a binomial random variable 
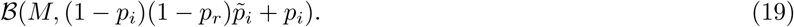

From (17), the left-hand-side of (16) is also binomially distributed 
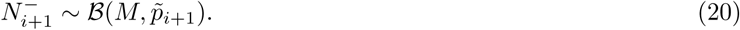

Equation (13) in the theorem results from matching the success probability parameter in (19) and (20). In summary, solving (13) based on some initial condition, i.e., 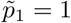 if all sites are occupied when the first action potential comes, provides the distribution for the number occupied sites 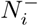 at time 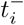 via (17). Using (14), (17), and the property of conditional binomials, the number of synaptic vesicles released is

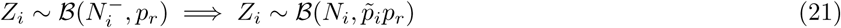

Two special cases resulting from Theorem 1 are

- *p_r_* = 1 (all occupied sites release vesicles), in which case 
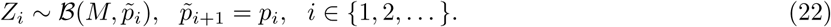
- *p_i_* = 1 (all empty sites get occupied before the next action potential), in which case 
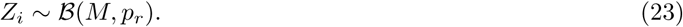

Next, we focus on computing the statistical moments of *Z_i_*.

## IV. MOMENTS OF THE NUMBER OF NEUROTRANSMITTERS RELEASED

Applying the formulas for the moments of a binomially distributed random variable in (3) to 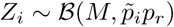 yields the following conditional moments 
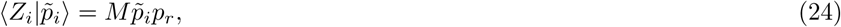

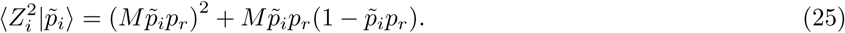

To uncondition (24), we first obtain the steady-state moments of 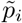 from (13). Taking the expected values on both sides of (13), and using the fact that *p_i_* are i.i.d. random variables drawn independently of 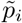, 
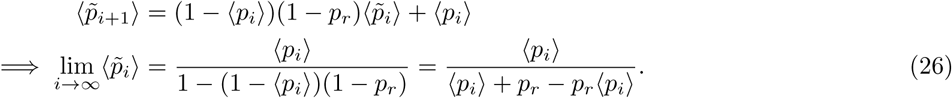

Similarly, squaring both sides in (13), taking the expected value and the limit *i* →∞ yields 
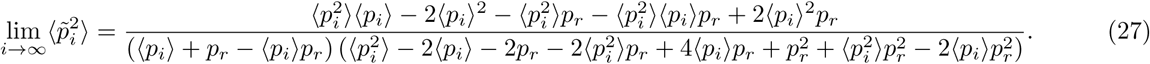

Now unconditioning (24) using the moments of 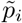, the average number of synaptic vesicles released per action potential at equilibrium is

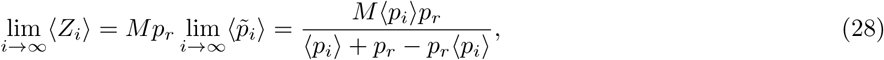
and using a similar approach, 
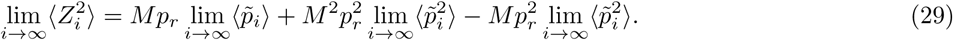

We now use these moments of *Z_i_* to investigate how stochasticity in the amount of neurotransmitter released can be regulated. In particular,

1. How does noise in *Z_i_* (quantified by its coefficient of variation squared) 
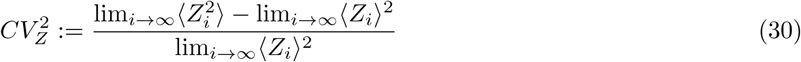
depend on noise in *T_i_* (action potential arrival times)?
2. Is there an optimal way to choose *M* (number of docking sites) and *p_r_* (vesicle release probability) so as to minimize 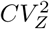?

To explore the first question, recall from (7) that *p_i_* is a monotonically increasing function of *T_i_*. Hence, increasing noise in the action potential timing can be modeled as increasing noise in *p_i_*. Fig. 2 plots 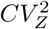 as a function of the coefficient of variation of *p_i_* for a fixed mean 〈*p_i_*〉. Our results reveal an interesting trade off: when the release probability is small, then 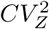 starts high but remains flat (Fig. 2). In contrast, when the release probability is close to one, then 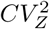 starts low but increases sharply.

**FIG.2.**
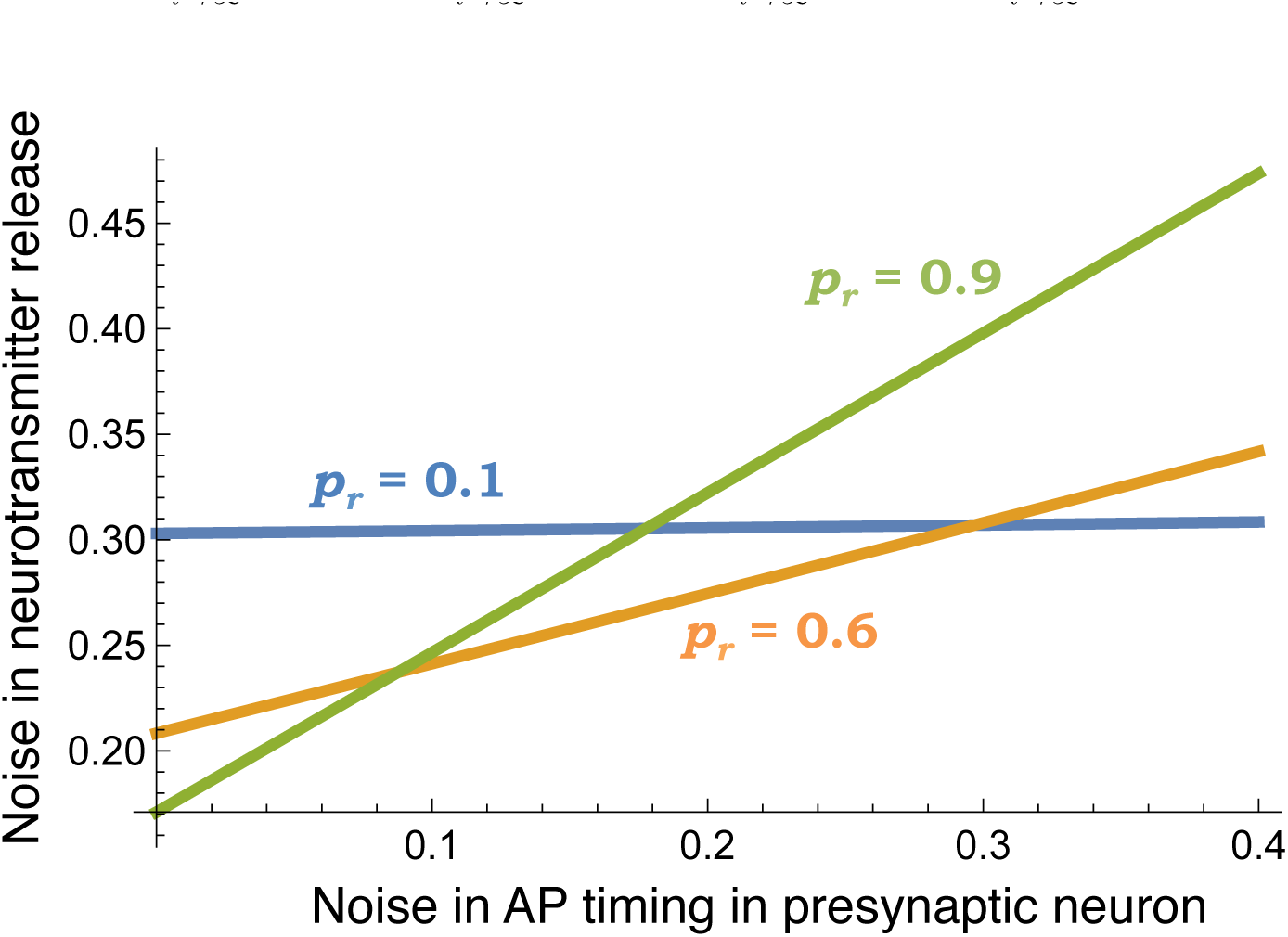
When noise in action potential (AP) timing is small (large) then a low (high) probability of release minimizes noise in neurotransmitter release. The coefficient of variation of *p_i_* is used as a proxy for noise in AP timing. 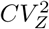 is plotted against the coefficient of variation of *p_i_* for 〈*p_i_*〉 = 0.75 and different values of *p_r_*. The number of docking sites is correspondingly changes as per (28) to keep the average of number of vesicles released lim*_i_*→∞〈Z*_i_*〉 = 3 fixed.

Fig. 3 plots 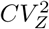 as a function of *p_r_* for deterministic arrivals (*T_i_* = 〈*T_i_*〉 with probability one), and Poisson arrivals (*T_i_* is exponentially distributed with mean 〈*T_i_*〉) of action potentials. The number of docking sites is correspondingly changes as per (28) to keep lim*_i_*→∞〈Z*_i_*〉 fixed. For deterministic arrivals, 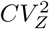 is always minimized by choosing *p_r_* = 1 (Fig. 3; top). Intriguingly, for Poisson arrivals, if the recovery probability 〈*p_i_*〉 is not large, then 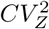 is minimized at an intermediate value of *p_r_* (Fig. 3; bottom). Note the qualitative shift in behavior when stochasticity is incorporated in *T_i_* (compare Fig. 3 bottom and top), and in some cases 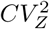 is minimized by choosing *p_r_* significantly smaller than one (see 〈*p_i_*〉 = 0.5 line in Fig. 3 bottom).

**FIG.3.**
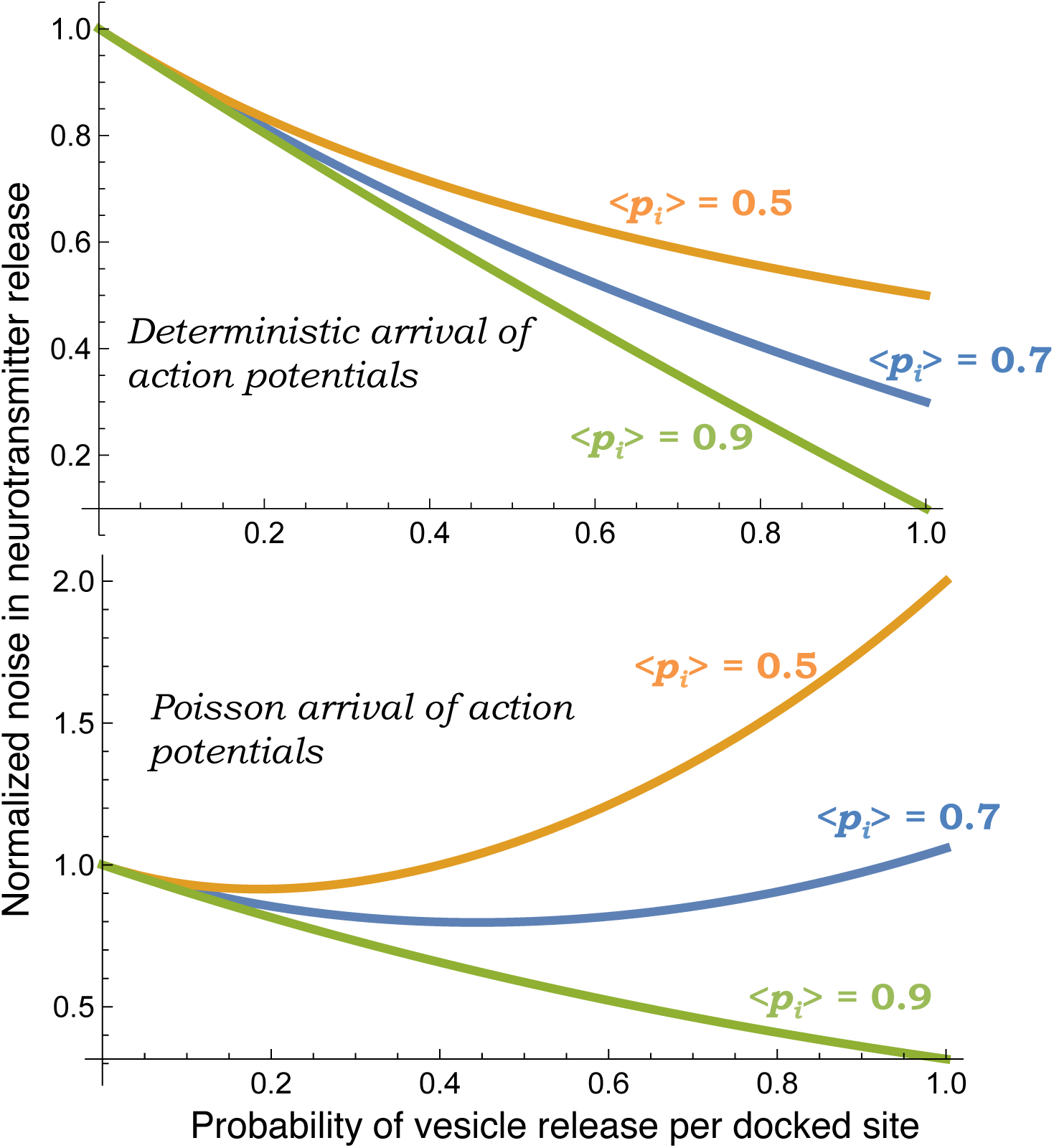
Noise in neurotransmitter release 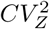 as a function of the vesicle release probability (*p_r_*) for deterministic (top) and Poisson (bottom) arrivals of action potential. In the former case, 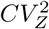 is always minimized at *p_r_* = 1 independent of the value of 〈*pi*〉 (top). In contrast, for Poisson arrivals, 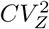 is minimized at a value of *p_r_* dependent on 〈*pi*〉 (bottom). More specifically, for high values of 〈*pi*〉, 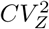 is again minimized at *p_r_* = 1, but for low values of 〈*pi*〉 the optima occurs at an intermediate value of *p_r_* (bottom). As in Fig. 2, the number of docking sites is correspondingly changes as per (28) to keep the average of number of vesicles released lim*_i_*→∞〈Z*_i_*〉 = 3 fixed. Noise levels are normalized to 1/lim*_i_*→∞〈Z*_i_*〉, the value of 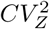 when *p_r_* = 0.

## V. ACTION POTENTIAL TIMING IN THE POSTSYNAPTIC NEURON

Having quantified the release of vesicles from the presynaptic neuron, we next focus on the stochastic dynamics of neurotransmitter counts in the synaptic gap (space between the cell membranes of the presynaptic and the postsynaptic neuron). Let *z*(*t*) denote the level of neurotransmitters in the synaptic gap, and its time evolution is characterized by production in bursts followed by exponential decay (Fig. 4). Whenever an action potential arrives at the presynaptic neuron, multiple vesicles are released causing a jump in *z*(*t*) 
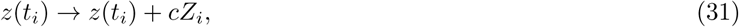
where random variable *Z_i_*, is the number of vesicles released and *c* is the amount of neurotransmitter per vesicle. In between action potentials, *z*(*t*) decreases as per a first-order decay process 
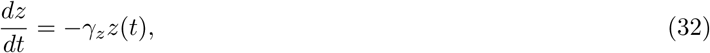
where *γ_z_* is the degradation rate of individual molecules. Given the average time between two action potentials (or burst events) 〈*T_i_*〉, the average number of neurotransmitters released per burst c〈*Z_i_*〉, and each molecule lives for an average time 1/*γ_z_* in the synaptic gap, we obtain the following steady-state mean levels from (28)^1^ 
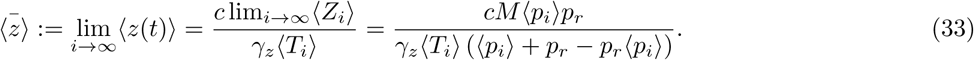

**FIG.4.**
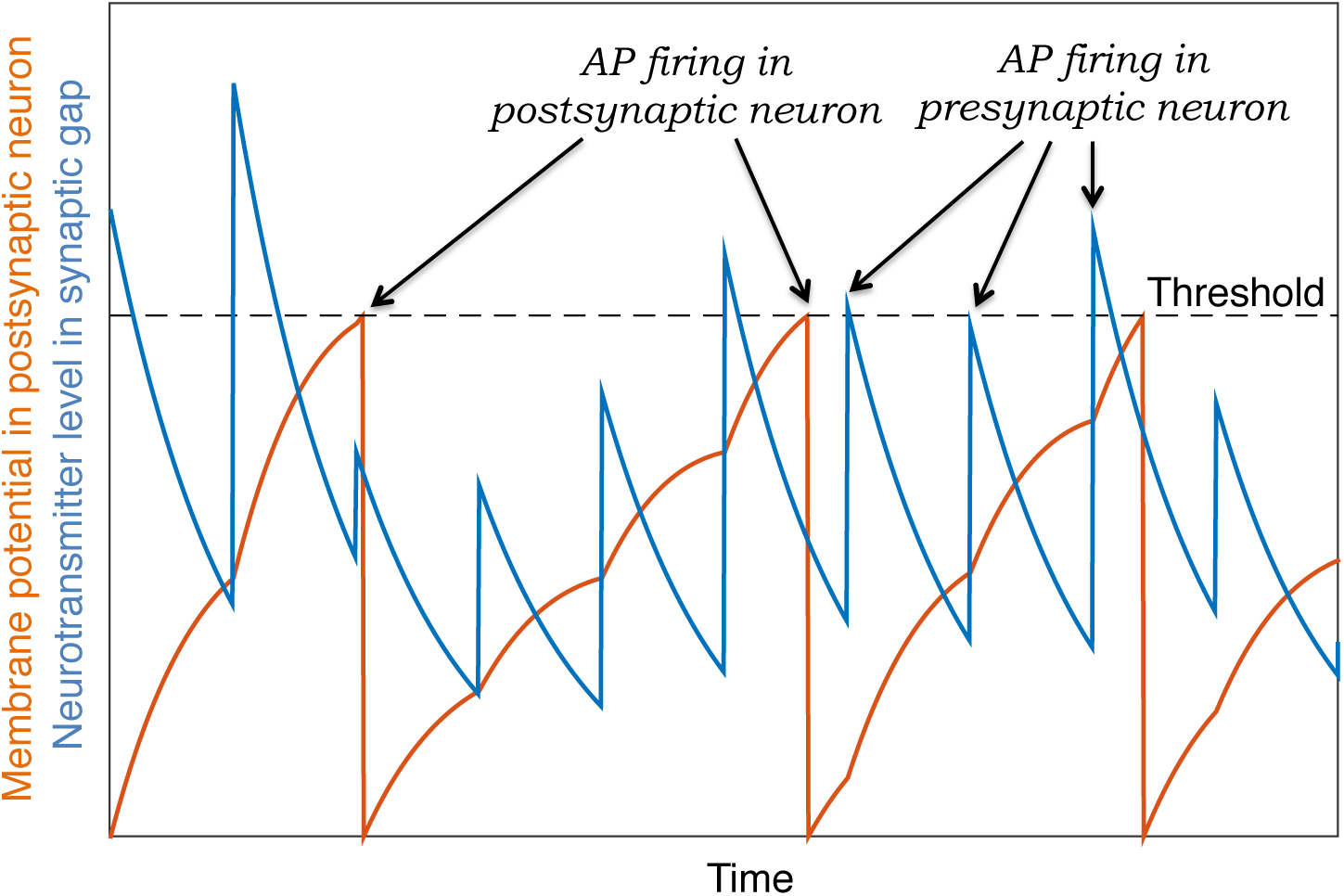
Generation of action potentials in the postsynaptic neuron based on the integrate-and-fire model. The neurotransmitter levels in the synaptic gap undergo bursty production and decay cycles, where burst events correspond to arrival of action potentials in the presynaptic neuron that lead to release of neurotransmitter-filled vesicles. The membrane potential evolves via (36), which can be interpreted as a “leaky integrator”. An action potential in the postsynaptic neuron is triggered when the potential hits a threshold for the first time. Immediately after this the membrane potential is reset to a lower value (membrane resting potential) and the process repeats.

Furthermore, recall that 
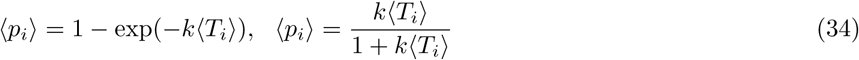
for deterministic and Poisson arrivals of action potentials, respectively. Substituting (34) in (33), one can see that the mean neurotransmitter level increases with increasing frequency of arrivals (1/〈*T_i_*〉), and at high frequencies

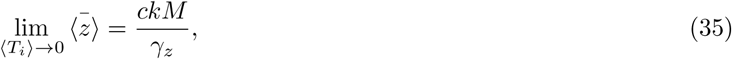
the system saturates to the same limit irrespective of deterministic or Poisson arrivals.

The stochastic process *z*(*t*) drives the linear dynamical system 
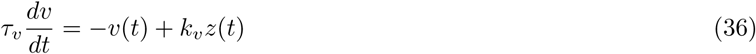
that determines when action potentials fire in the postsynaptic neuron. Model (36) is commonly referred to as the *integrate-and-fire model* [18], and here *v*(*t*) is the membrane potential of the postsynaptic neuron, *τ_υ_* is the membrane time constant and *k_v_* is a positive constant. An action potential in the postsynaptic neuron is triggered when *v*(*t*) hits a threshold *v_th_* for the first time. At this time, *v*(*t*) is reset to zero and the process starts anew. We use *v* = 0 to represent the membrane resting potential that is typically −70 *mV*. In essence, the time to the next action potential in the postsynaptic neuron is the first-passage time

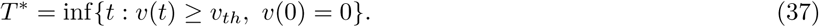

Note that if fluctuations in *z*(*t*) are long lived (for example, low turnover rate of neurotransmitters in the synaptic gap), then the timing of successive action potentials in the postsynaptic neuron will be correlated.

Since fluctuations in *z*(*t*) are at steady-state, the mean membrane potential increases as 
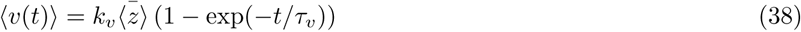
starting from *υ*(0) = 0. Assuming 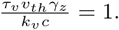 and small fluctuations in *T\**, the mean first-passage time can be approximated using 
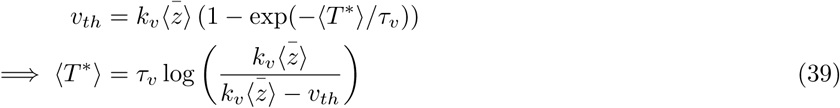

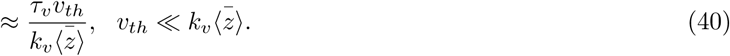

Using (33)-(34) in (40), Fig. 5 plots the average frequency of action potentials in the postsynaptic neuron (1 */T\**) as a function of the input frequency (1 */*〈*T_i_*〉) and shows a linear increase followed by saturation. The saturation limit is obtained by substituting (35) in (40), and interestingly, this limit is dependent on *M*, but completely independent of *p_r_*. Thus, at sufficiently high input frequencies of action potentials, changing *p_r_* will not affect the action potential timing in the postsynaptic neuron.

**FIG.5.**
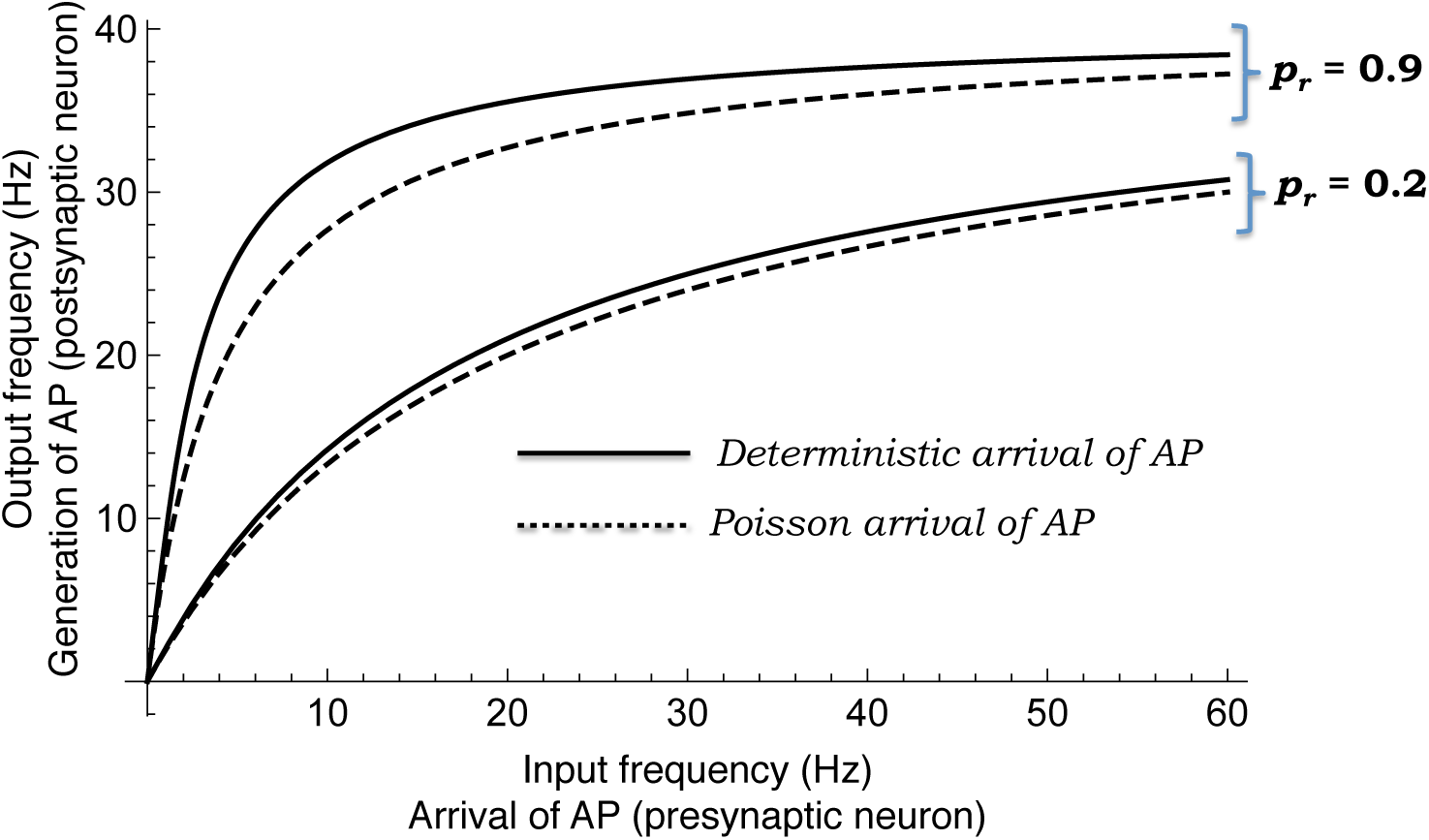
As the action potential (AP) frequency increases in the presynaptic neuron, the action potential frequency (output frequency) in the postsynaptic neuron increases and saturates. A higher probability of vesicle release causes a sharper increase, but the same saturation limit that is determined by substituting (35) in (40). Moreover, deterministic AP arrivals lead to higher output frequency compared to Poisson arrivals for the same 〈*T_i_*〉. Other parameters taken as *M* = 10, *k* = 4 *sec^-1^* and the ratio 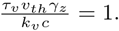

## VI. NOISE IN ACTION POTENTIAL TIMING

Having determined the mean time to the next action potential, the focus now is on the noise properties of *T\**. We assume that the noise in the first-passage time *T\** is sufficiently small, that it is simply proportional (or monotonically related) to the noise in *υ*(*t*) computed at the mean first-passage time *t* = 〈*T*\*〉. This assumption naturally motives the question: Can we derive closed-form expressions for temporal dynamics of *v*(*t*) noise levels?

Recall from the previou section that random processes *z*(*t*) and *υ*(*t*) are defined through the stochastic hybrid system (31), (33) and (36). To aid moment computations described below, we assume Poisson arrivals of action potentials and

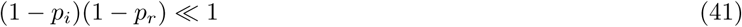
in (13) which makes 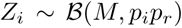 an i.i.d random variable. Referring interested readers to [19–21] for details on moment dynamics for stochastic hybrid systems, the time evolution of all the first and second-order moment of *z*(*t*), *υ*(*t*) is obtained as

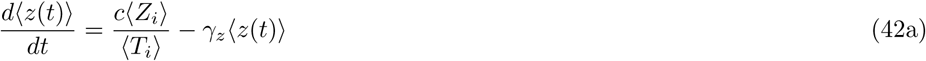

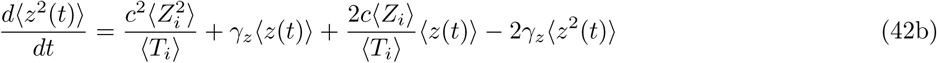

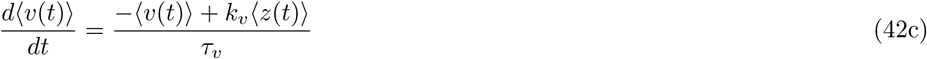

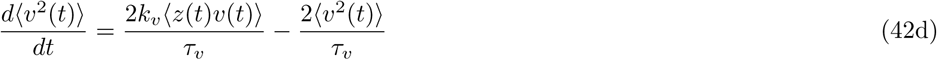

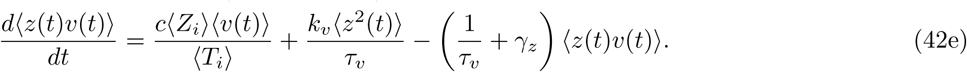

Based on the first two equation in (42), the steady-state moments of *z*(*t*) are

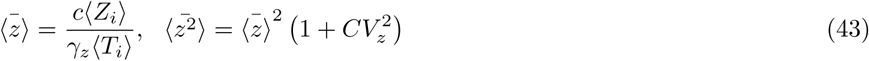

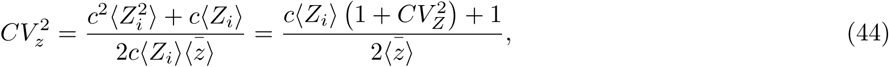
where 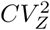 is the noise in the neurotransmitter release process defined in (30) ^2^. Since *v*(0) = 0 just after an action potential firing, and fluctuations in *z*(*t*) are at steady-state, solving the linear dynamical system (42) with the following initial conditions 
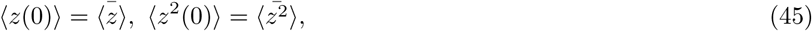

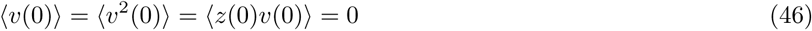
yields the following noise levels in *υ*(*t*) (quantified by its coefficient of variation squared)

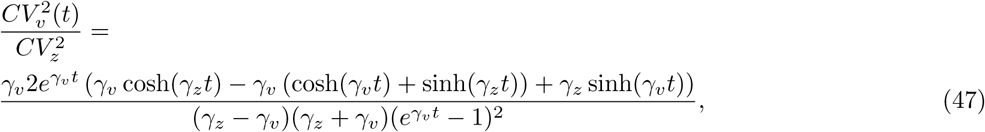
where 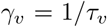 and cosh (sinh) are hyperbolic cosine (sine) functions. Based on our earlier assumption, and given (47), noise in the first-passage time *T*\* is 
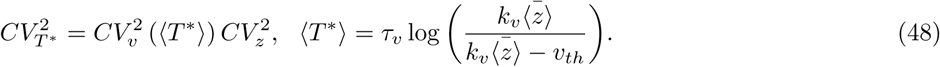

As one would expect, random fluctuations in *T*\* are connected to noise in the neurotransmitter counts 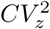, which in turn is related to noise in Z*_i_* via (43). Thus, mechanisms to buffer noise in *Z_i_* illustrated in Figs. 2 & 3 are critical for achieving precise action potential firing times. Fig. 6 plots 〈*T*\*〉 and 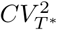 as a function of the firing threshold *υ^th^* - while the average time to the next action potential increases, the noise in *T*^*^ decreases with increasing threshold. Furthermore, it can be seen from (48) that 
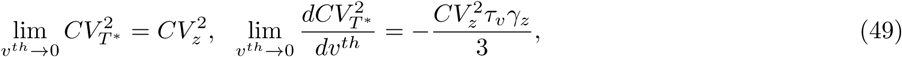
and hence the decrease in noise levels in Fig. 6 is sharper for higher membrane time constant *τ_ν_*, or faster turnover rate of neurotransmitters *γ_z_*.

**FIG. 6.**
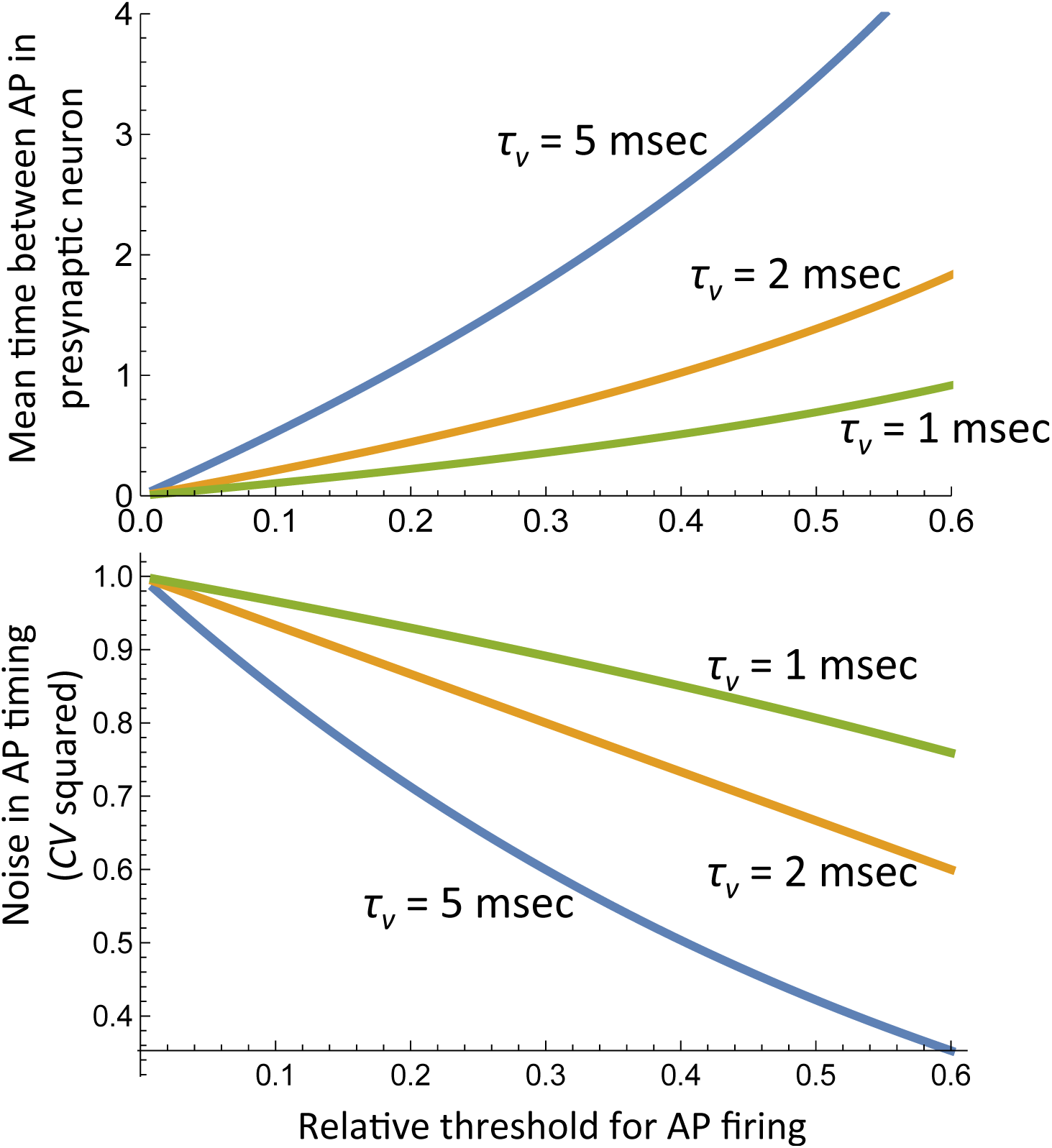
The mean time to the next action potential (top) and noise in timing 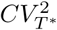 (bottom) as a function of the relative firing threshold 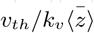 for different value of *τ_ν_*. While the mean time 〈*T*\*〉 increases, the noise 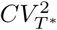 decreases with increasing *υ_th_*. For this plot *γ_z_* = 1 *msec*. The noise levels are normalized by their value for *υ_th_* = 0.

## VII. DISCUSSION

We have performed a systematic analysis of noise mechanisms in synaptic transmission between two neuron pairs. Our key mathematical contributions are

- Quantifying the statistics of neurotransmitter release, each time the presynaptic neuron is stimulated by an action potential (Theorem 1).
- Characterizing fluctuations in the number of neurotransmitter molecules in the synaptic gap.
- Derivation of approximate formulas for the timing of action potentials in the postsynaptic neuron based on the integrate-and-fire modeling framework.

These results have led to some intriguing biological insights that are summarized below. Counter-intuitively, when action potentials randomly arrive at the presynaptic neuron, having an optimal probability of releases *p_r_* minimizes the noise in the number of neurotransmitters released (Figs. 2 & 3). Furthermore, in some cases deterministically releasing vesicles based on *p_r_* = 1 would considerably amplify noise (Fig. 3; bottom). Our results further show that high frequencies of action potentials in the presynaptic neuron lead to saturation in the neurotransmitter count, and the action potential frequency in the postsynaptic neuron (Fig. 5). Interestingly, this saturation limit depends on the number of docking sites, but completely independent of *p_r_*. Finally, our model analysis reveals that increasing the firing threshold in integrate-and-fire models can buffer noise and enhance precision in action potential timing (Fig. 6).

In recent work, we have developed exact analytical results for the first-passage time in stochastic models of bursty gene expression [24–26]. Motivated by this literature, an important direction of future research is to perform a mathematically rigorous derivation of the first-passage time *T^*^*, and explore how the results in Fig. 6 change when the magnitude of fluctuations in *T^*^* are large, or when threshold crossings are noise induced. Other directions of future work include:

- Expand the current model to consider multiple synapses (both inhibitory and excitatory).
- Investigate how negative feedback in the form of an autapse (synapse from a neuron onto itself) can enhance noise buffering in synaptic transmission [27, 28].
- While the current work assumes biophysical parameters, such as, the number of docking sites and release probability to be constant, experimental work points to dynamic regulation of these parameters [29–31]. It will be interesting to investigate the effects of such dynamic regulation through local feedbacks on synaptic transmission.

## ACKNOWLEDGMENT

AS is supported by the National Science Foundation Grant DMS-1312926, University of Delaware Research Foundation (UDRF) and Oak Ridge Associated Universities (ORAU).

## Appendix A

Here we prove that given (18), the right-hand-side of (16) is binomially distributed as

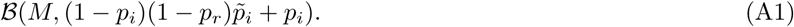

The MGF of the right-hand-side of (16) conditioned on 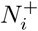 is given by

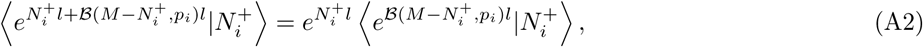

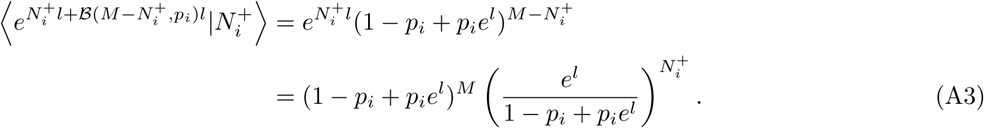

Now unconditioning (A3) with respect to 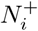 yields

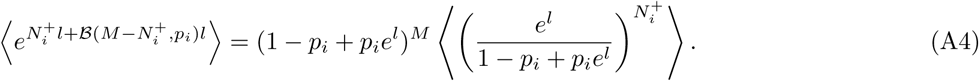

Since 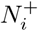 is also a Binomial random variable, we have from (4) and (18)

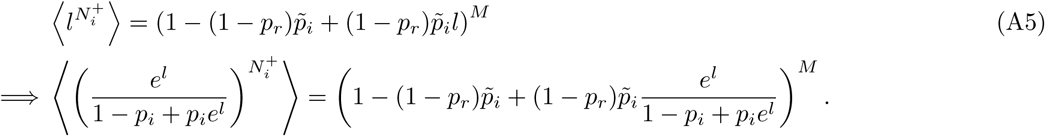

Simplifying (A4) using (A6)

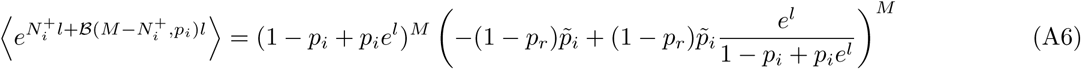

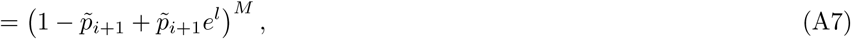
where 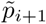 is given by (13) and (A7) is the MGF of a binomial random variable 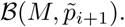.

## Footnotes

1 If we think molecules in the synaptic gap as customers in a queue, this result, can be derived based on Little’s law for a *G/M/∞* queue, where customers arrives in batches based on some general inter-burst, time distribution *T_i_* [17].

2 Results in (44) are analogous to protein noise levels in bursty models of gene expression [19, 22, 23], with the only exception that *Z_i_* is binomially distributed, while protein burst sizes typically follow geometric distributions

## References

[1] R. S. Zucker and W. G. Regehr, “Short-term synaptic plasticity,” Annual Review of Physiology, vol. 64, pp. 355–405, 2002.

[2] M. Tsodyks and H. Markram, “The neural code between neocortical pyramidal neurons depends on neurotransmitter releaseprobability,” Proceedings of the National Academy of Sciences, vol. 94, pp. 719–723, 1997.

[3] L. F. Abbott, J. A. Varela, K. Sen, and S. B. Nelson, “Synaptic depression and cortical gain control,” Science, vol. 275, pp. 221–224, 1997.

[4] M. Tsodyks, K. Pawelzik, and H. Markram, “Neural Networks with Dynamic Synapses,” Neural Computation, vol. 10, pp. 821–835, 1998.

[5] H. Markram, Y. Wang, and M. Tsodyks, “Differential signaling via the same axon of neocortical pyramidal neurons,” Proceedings of the National Academy of Sciences, vol. 95, pp. 5323–5328, 1998.

[6] W. Senn, H. Markram, and M. Tsodyks, “An Algorithm for Modifying Neurotransmitter Release Probability Based on Pre- and Postsynaptic Spike Timing,” Neural Computation, vol. 13, pp. 35–67, 2001.

[7] C. Pulido, F. Trigo, I. Llano, and A. Marty, “Vesicular release statistics and unitary postsynaptic current at single {GABAergic} synapses”, Neuron, vol. 85, pp. 159–172, 2015.

[8] G. Malagon, T. Miki, I. Llano, E. Neher, and A. Marty, “Counting vesicular release events reveals binomial release statistics at single glutamatergic synapses”, Journal of Neuroscience, vol. 36, pp. 4010–4025, 2016.

[9] A. A. Faisal, L. P. J. Selen, and D. M. Wolpert, “Noise in the nervous system”, Nature Reviews Neuroscience, vol. 9, pp. 292–303, 2008.

[10] F. S. Chance, S. B. Nelson, and L. F. Abbott, “Synaptic depression and the temporal response characteristics of v1 cells”,; Journal of Neuroscience, vol. 18, pp. 4785–4799, 1998.

[11] R. Rosenbaum, J. E. Rubin, and B. Doiron, “Short-term synaptic depression and stochastic vesicle dynamics reduce and shape neuronal correlations”, Journal of Neurophysiology, vol. 109, pp. 475–484, 2013.

[12] R. Rosenbaum, J. Rubin, and B. Doiron, “Short term synaptic depression imposes a frequency dependent filter on synaptic information transfer”, PLOS Computational Biology, vol. 8, p. e1002557, 2012.

[13] M. S. Goldman, “Enhancement of Information Transmission Efficiency by Synaptic Failures”,; Neural Computation, vol. 16, pp. 1137–1162, 2004.

[14] E. Schneidman, B. Freedman, and I. Segev, “Ion Channel Stochasticity May Be Critical in Determining the Reliability and Precision of Spike Timing”, Neural Computation, vol. 10, pp. 1679–1703, 1998.

[15] C. Zhang and C. S. Peskin, “Improved signaling as a result of randomness in synaptic vesicle release”, Proceedings of the National Academy of Sciences, vol. 112, pp. 14 954–14 959, 2015.

[16] A. Arleo, T. Nieus, M. Bezzi, A. D’Errico, E. D’Angelo, and O. J.-M. D. Coenen, “How Synaptic Release Probability Shapes Neuronal Transmission: Information-Theoretic Analysis in a Cerebellar Granule Cell”, Neural Computation, vol. 22, 2010.

[17] S. Asmussen, Applied Probability and Queues. Springer, 2003.

[18] C. Koch and I. Segev, Methods in Neuronal Modeling: From Ions to Networks, ser. A Bradford book. MIT Press, 1998.

[19] A. Singh, “Negative feedback through mRNA provides the best control of gene-expression noise”, IEEE Transactions on Nanobioscience, vol. 10, pp. 194–200, 2011.

[20] J. P. Hespanha and A. Singh, “Stochastic models for chemically reacting systems using polynomial stochastic hybrid systems”, International Journal of Robust and Nonlinear Control, vol. 15, pp. 669–689, 2005.

[21] A. Singh and J. P. Hespanha, “Stochastic hybrid systems for studying biochemical processes”, Philosophical Transactions of the Royal Society A, vol. 368, pp. 4995–5011, 2010.

[22] A. Singh, “Transient changes in intercellular protein variability identify sources of noise in gene expression”, Biophysical Journal, vol. 107, pp. 2214–2220, 2014.

[23] A. Singh and J. P. Hespanha, “Optimal feedback strength for noise suppression in autoregulatory gene networks”,; Biophysical Journal, vol. 96, pp. 4013–4023, 2009.

[24] K. R. Ghusinga, J. J. Dennehy, and A. Singh, “First-passage time approach to controlling noise in the timing of intracellular events”, Proceedings of the National Academy of Sciences, vol. 114, pp. 693–698, 2017.

[25] K. R. Ghusinga, C. A. Vargas-Garcia, and A. Singh, “A mechanistic stochastic framework for regulating bacterial cell division”,; Scientific Reports, p. 30229, 2016.

[26] K. R. Ghusinga and A. Singh, “First-passage time calculations for a gene expression model,” Proc. of the 53rd IEEE Conf. on Decision and Control, Los Angeles, CA, pp. 3047–3052, 2014.

[27] A. Bacci and J. R. Huguenard, “Enhancement of spike-timing precision by autaptic transmission in neocortical inhibitory interneurons”,; Neuron, vol. 49, pp. 119–130, 2006.

[28] D. Guo, S. Wu, M. Chen, M. Perc, Y. Zhang, J. Ma, Y. Cui, P. Xu, Y. Xia, and D. Yao, “Regulation of Irregular Neuronal Firing by Autaptic Transmission”, Scientific Reports, vol. 6, p. 26096, 2016.

[29] T. Ngodup, J. A. Goetz, B. C. McGuire, W. Sun, A. M. Lauer, and M. A. Xu-Friedman, “Activity-dependent, homeostatic regulation of neurotransmitter release from auditory nerve fibers”, Proceedings of the National Academy of Sciences, vol. 112, pp. 6479–6484, 2015.

[30] A. G. Millar, H. Bradacs, M. P. Charlton, and H. L. Atwood, “Inverse relationship between release probability and readily releasable vesicles in depressing and facilitating synapses”, Journal of Neuroscience, vol. 22, pp. 9661–9667, 2002.

[31] T. Branco and K. Staras, “The probability of neurotransmitter release: variability and feedback control at single synapses”, Nature Reviews Neuroscienc, vol. 10, pp. 373–383, 2009.

